# Role of the ubiquitin ligase ITCH in clathrin-mediated endocytosis of the epidermal growth factor receptor

**DOI:** 10.1101/2021.02.15.431259

**Authors:** Riham Ayoubi, Peter S. McPherson, Annie Angers

## Abstract

Once activated by ligand, epidermal growth factor receptor (EGFR) is endocytosed in clathrin-coated pits. ITCH is an E3 ubiquitin ligase that interacts with and ubiquitinates several proteins involved in clathrin-mediated endocytosis (CME) including endophilin. To further investigate the function of ITCH in EGFR endocytosis, the internalization of fluorescent EGF was measured in ITCH^-/-^ HeLa cells. In the absence of ITCH, there was a significant decrease in the CME of EGF. Rescue experiments using wild-type ITCH confirmed the importance of the protein for normal EGF uptake. ITCH point mutations that disrupt the interaction of ITCH with endophilin failed to rescue the defects in EGFR uptake, as did a non-catalytic form of ITCH. ITCH^-/-^ cells also displayed a delay in the rate of phospho-EGFR degradation as well as prolonged ERK1/2 signaling. Our study uncovers a pathway regulating EGFR trafficking and reveals for the first time that the protein ITCH is required for CME of EGFR.

## Introduction

Uptake of extracellular growth factors occurs through ligand-dependent endocytosis of cell surface receptors. Upon ligand binding, the plasma membrane invaginates to form clathrin-coated pits (CCPs) that harbor receptors such as the epidermal growth factor receptor (EGFR), and these CCPs subsequently pinch off and deliver the ligand-receptor complex to endosomes. During this process, EGFR transduces signals important for cell growth, differentiation, proliferation, and motility. Once the clathrin-dependent route is saturated, clathrin-independent endocytosis takes place allowing fast degradation of the receptor in lysosomes (1,2).

Ubiquitination is a post-translational modification that can regulate receptor trafficking, sorting, and downregulation (3). Following EGFR dimerization, auto-phosphorylation at tyrosine 1045 (pY1045) allows the ubiquitin ligase CBL to bind and ubiquitinate the receptor, a signal that targets EGFR to degradation in lysosomes (4–6). Interestingly, CME is also regulated by ubiquitination of endocytic adaptors. For example, CBL itself is a substrate for another ubiquitin ligase, ITCH (also known as AIP4, atrophin-interacting protein 4), which regulates CBL levels (7).

ITCH is a ubiquitin ligase that contains an N-terminal C2 domain, four WW domains and a homologue of the E6-AP carboxyl terminus (HECT) catalytic domain. ITCH typically uses its WW domains to interact with PPxY motifs of its substrates, such as CBL. However, ITCH also contains a proline rich region (PRR) that recognizes several of the SH3 domain-containing proteins involved in CME. These include endophilin, amphiphysin, PACSIN, Grb2 (8,9), and SNX9 (10). We previously tested the affinity of the ITCH PRR for these different SH3 domain proteins and discovered that the affinity for endophilin was the highest amongst those tested (9). Here, we demonstrate that an ITCH protein with mutations in arginine 252,255, and 258 is unable to bind endophilin.

Endophilins are N-BAR proteins that participate in the internalization of EGFR by sensing and driving membrane curvature, while acting as a scaffold to recruit other crucial endocytic proteins (11,12). For example, a CBL/CIN85/endophilin complex mediates downregulation of EGFR following receptor activation (13). The interactions of ITCH with CBL, endophilin and other SH3-domain containing proteins suggests a role for ITCH in the internalization process of the EGF receptor. Therefore, we examined whether loss-of-function of ITCH affects the endocytosis of stimulated EGFR. We used CRISPR/Cas9 to knockout (KO) ITCH in HeLa cells and found that it is required for ligand-bound EGFR but not the transferrin receptor endocytosis. Importantly, while the overexpression of wild-type ITCH successfully restores the level of internalized EGF/EGFR, neither a ligase-dead mutant nor a PRR mutant unable to bind endophilin rescues the phenotype. The present study provides an example of a HECT domain ubiquitin ligase functioning as an important regulator of EGFR endocytosis, further enhancing the importance and complexity of both EGFR signaling and the ubiquitination process as a regulator of endocytic mechanisms.

## Experimental procedures

### Cell culture and transfection

HEK-293T, HeLa and COS-7 cells were grown in high-glucose DMEM (GE Healthcare) supplemented with 10% bovine calf serum (GE Healthcare), penicillin (Invitrogen, 100 units/mL), streptomycin (Wisent, 100 μg/ml) and 2 mM L-glutamate in an incubator set at 5% CO_2_ and 37°C. HEK-293T cells were transfected using the calcium/phosphate method and HeLa cells using jetPRIME^®^ transfection reagent (Polyplus) according to the manufacturer’s instructions.

### Antibodies

Purified Mouse Anti-Itch/ITCH from BD Biosciences (Catalog #611198), Rabbit polyclonal anti-c-CBL from Cell Signaling (Catalog #2747), Rabbit polyclonal anti-endophilinA2 from Abcam (Catalog #ab130600), Rabbit monoclonal EGFR from Cell signaling (D38B1), Mouse mAb antip44/42 MAPK (Erk1/2) (Catalog #9107), Rabbit polyclonal anti-Phospho-EGF Receptor (Tyr1045) (Catalog #2237), Phospho-EGF Receptor (Tyr1148) (Catalog #4404) and Rabbit monoclonal Phospho-EGF Receptor D7A5 (Tyr1068) (Catalog #3777) from Cell signaling, Rabbit polyclonal anti-GFP from Invitrogen (Catalog #A-6455) and Anti-β Actin from Millipore (Catalog #MAB1501) were used for western blot. Mouse monoclonal anti-extracellular EGFR from Abcam (ab30) was used for immunofluorescence.

### Mutagenesis

All GFP-ITCH Full mutants were generated using QuikChange II site-directed mutagenesis kit (Agilent Technologies) according to the manufacturer’s instructions and using primers designed to introduce point mutations by changing arginine to glutamic acid (Table 1).

**Table 1.**
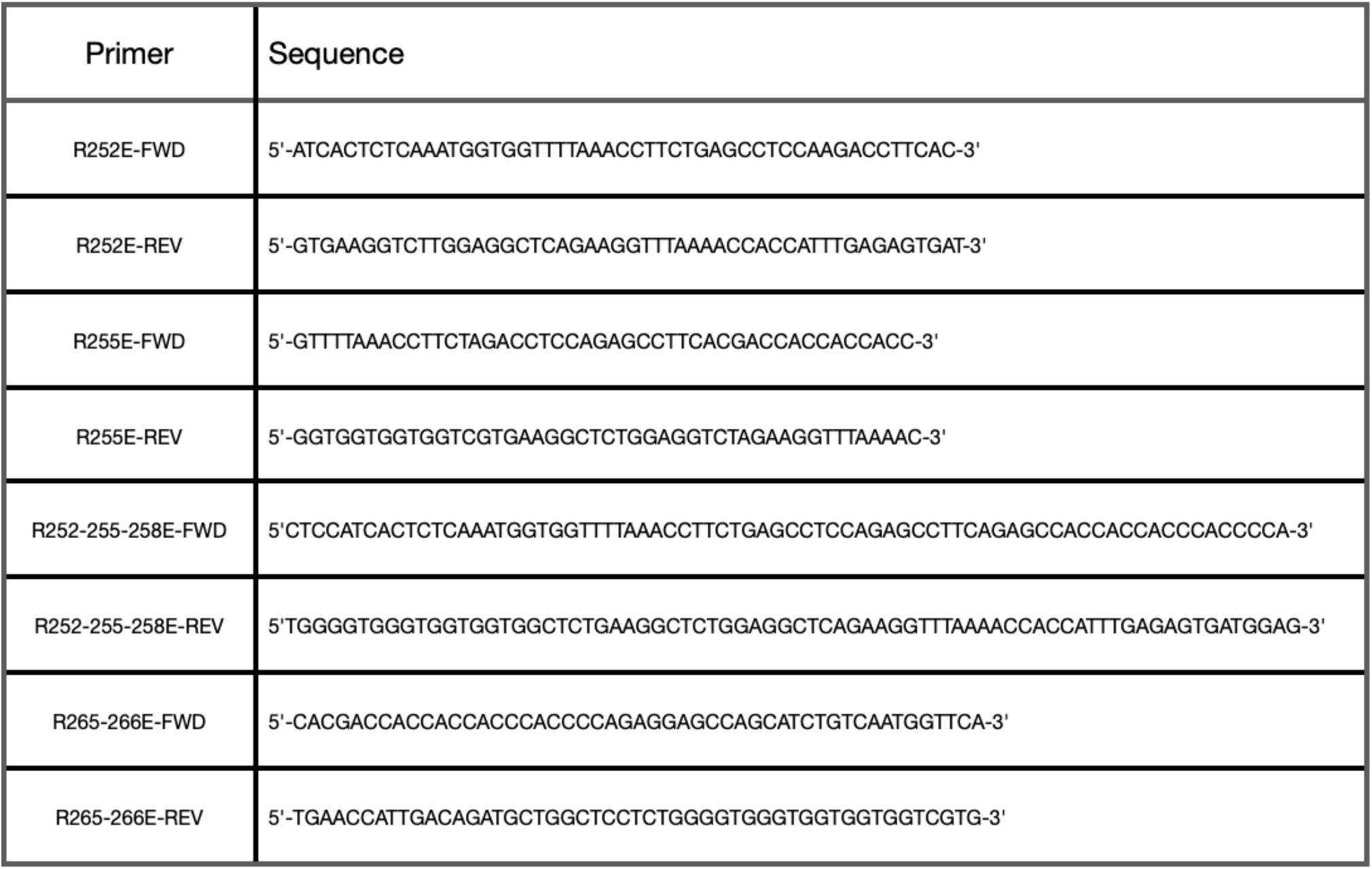
Primers used for ITCH mutagenesis.

### CRISPR/Cas9 KO

CRISPR/Cas9-mediated ITCH KO was performed as described (14). Briefly, for HeLa cells, guide RNAs were designed to target a unique sequence in the ITCH human genomic DNA using the Benchline online tool. Two guide RNAs positioned 100 bp apart were selected (Table 2). They were cloned individually into pSpCas9(BB)-2A-Puro (PX459) V2.0 vector and both vectors were transfected into the cells. For COS-7 cells, one guide RNA was used (Table 2). At 24h posttransfection, the cells were treated with puromycin at a concentration determined by a puromycin kill curve on parent cells. After selection, cells were diluted to obtain a single cell colony. Single colonies were expanded, DNA was extracted using QuickExtract genomic DNA extraction solution (Epicentre Biotechnologies) and then screened by PCR amplification of the region of interest. The knockout was verified by sequencing the DNA modification and by immunoblot.

**Table 2.**
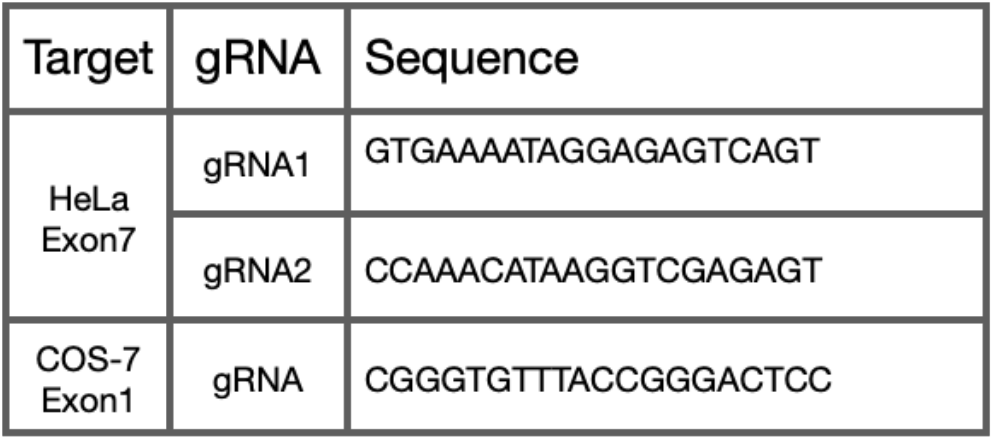
Guide RNAs used for ITCH CRISPR/Cas9 knockout.

### Endocytosis assay and rescue experiments

Cells plated on coverslips were serum-starved in DMEM with 2 mM L-glutamate for 2 h at 37°C. Alexa Fluor 647-EGF (Invitrogen; Catalog #E35351) diluted in cold serum-free media was then added, and cells were kept at 4°C in the dark for 1 h. EGF-free media at 37°C was added and cells were then allowed to internalize the bound EGF for 20 min at 37°C. Cells were then gently acid washed (0.2M acetic acid/0.4M NaCl) for 30 sec to remove any EGF bound to the extracellular membrane and fixed in 4% Paraformaldehyde.

For rescue experiments, parent HeLa cells or ITCH KO cells were transfected with empty pEGFP-C2 vector and ITCH KO cells were transfected with the indicated GFP-ITCH constructs. At 24 h post transfection, endocytosis assays were performed using 2 ng/ml Alexa Fluor-647 EGF. A similar endocytosis assay was used to observe transferrin uptake using 100 μg/mL of Alexa Fluor-647 (Catalog #T23366) or 568 (Catalog #T23365) conjugated transferrin (Invitrogen).

Mosaic experiments were performed to directly compare parent and ITCH^-/-^ cells in the same microscopic field. In these experiments, parent and ITCH KO cells were transfected with pEGFP-C2 and mCherry, respectively. At 24 h post transfection, cells were detached and mixed at a 1:1 ratio before seeding on coverslips for another 24 h. Endocytosis assays were then performed at 2 ng/ml Alexa Fluor-647 EGF.

### Pull-down assays

All GST fusion proteins were expressed in *Escherichia coli* DH5α and purified on Glutathione-Sepharose beads in HEPES buffer supplemented with a protease inhibitor cocktail. HEK-293T cells were transfected with different GFP-ITCH constructs. The next day, cells were washed 3X with PBS, lysed and incubated for 20 min at 4°C with 20 mM HEPES buffer with protease inhibitors supplemented with Triton-X100 at 1% final concentration. Lysates were centrifuged at 4°C for 15 min at 15000 rpm using a microcentrifuge. Supernatants were then incubated with the above-mentioned fusion proteins overnight at 4°C followed by elution in 1x SDS-PAGE sample buffer. All eluted samples were then run on SDS-PAGE denaturing gels followed by immunoblotting.

### Imaging and statistical analysis

Cells were imaged on a Leica TCS-SP8 confocal microscope using a 63× oil immersion objective. Images were processed using the LasX software. Internalized AF647-EGF was quantified in 60 to 90 cells for each condition in three independent experiments by drawing a constant area box around each cell using ImageJ. Corrected total cell fluorescence was compared between groups using R-studio software (R Core Team, 2016). The normality and homogeneity of data were verified using the Shapiro-Wilk test for the former and Levene test for the latter. Depending on the specific biological question, statistical analyses were carried out using a two-way ANOVA, one-way ANOVA (followed by Bonferroni’s post hoc comparison) or Student’s t test. P-values < 0.001 were considered significant.

### Degradation chase experiments

Serum-starvation medium was supplemented with 100 μg/ml cycloheximide to block new protein synthesis. Cells were deprived of serum for 2h and stimulated with 2 ng/ml of recombinant human EGF (Peprotech; Catalog #AF-100-15) for 2, 5, 10, 20, 30, 60, 120, 240 min. Cells were lysed in 20 mM HEPES buffer, 1% Triton-X-100 with protease and phosphatase inhibitors and centrifuged for 15min at 15000 rpm. Total cell lysates were separated on denaturing SDS-PAGE and immunoblotted with the indicated antibodies.

## Results

### KO of ITCH causes a deficit in CME of EGFR

Following binding to EGF, EGFR is internalized to be degraded in lysosomes (15). Since multiple SH3-domain containing proteins involved in endocytosis are known to bind the ubiquitin ligase ITCH (9,10,16,17), we hypothesized that the absence of ITCH would influence the endocytosis of EGFR. To test this hypothesis, we generated ITCH KO cells using CRISPR/Cas9. At low concentrations, at or below 2 ng/ml, EGF triggers CME of the EGFR, but at concentrations above ~ 2-5 ng/ml EGF, EGFR saturates the CME pathway and additionally enters cells through clathrin-independent endocytosis (6,18,19). We thus tested EGFR internalization at various concentrations of EGF (2, 5, 10 and 20 ng/ml). The cells were briefly acid washed before fixation so that only internalized fluorescent EGF was visible. We quantified endocytosed EGF and compared the total fluorescence of EGF-positive puncta between ITCH KO and parental HeLa cells. Upon incubation of the cells with 2 ng/ml of fluorescent EGF at 37°C for 20 min, a significant decrease in the amount of internalized EGF was observed in ITCH KO cells compared to parental controls (Fig. 1A/C). This phenotype was gradually less visible when higher concentrations of EGF were used, such that by 5 ng/ml the difference seemed less obvious and at 10 or 20 ng/ml there was seemingly no difference between KO and parental cells (Fig. 1A). Statistical analysis revealed significant differences only at 2 ng/ml (Fig. 1C and S1). Identical results were seen in COS-7 cells (Fig. 1B/D).

The alterations in EGFR endocytosis occurred despite the fact there were no changes in the levels of the ITCH substrates Endophilin-A2 or c-CBL in the ITCH KO cells. Moreover, endogenous levels of total EGFR remained unchanged in control and ITCH KO cells (Fig. S3A) and there was no change in the level of EGFR at the cell surface (Fig. S3B/C).

**Figure 1.**
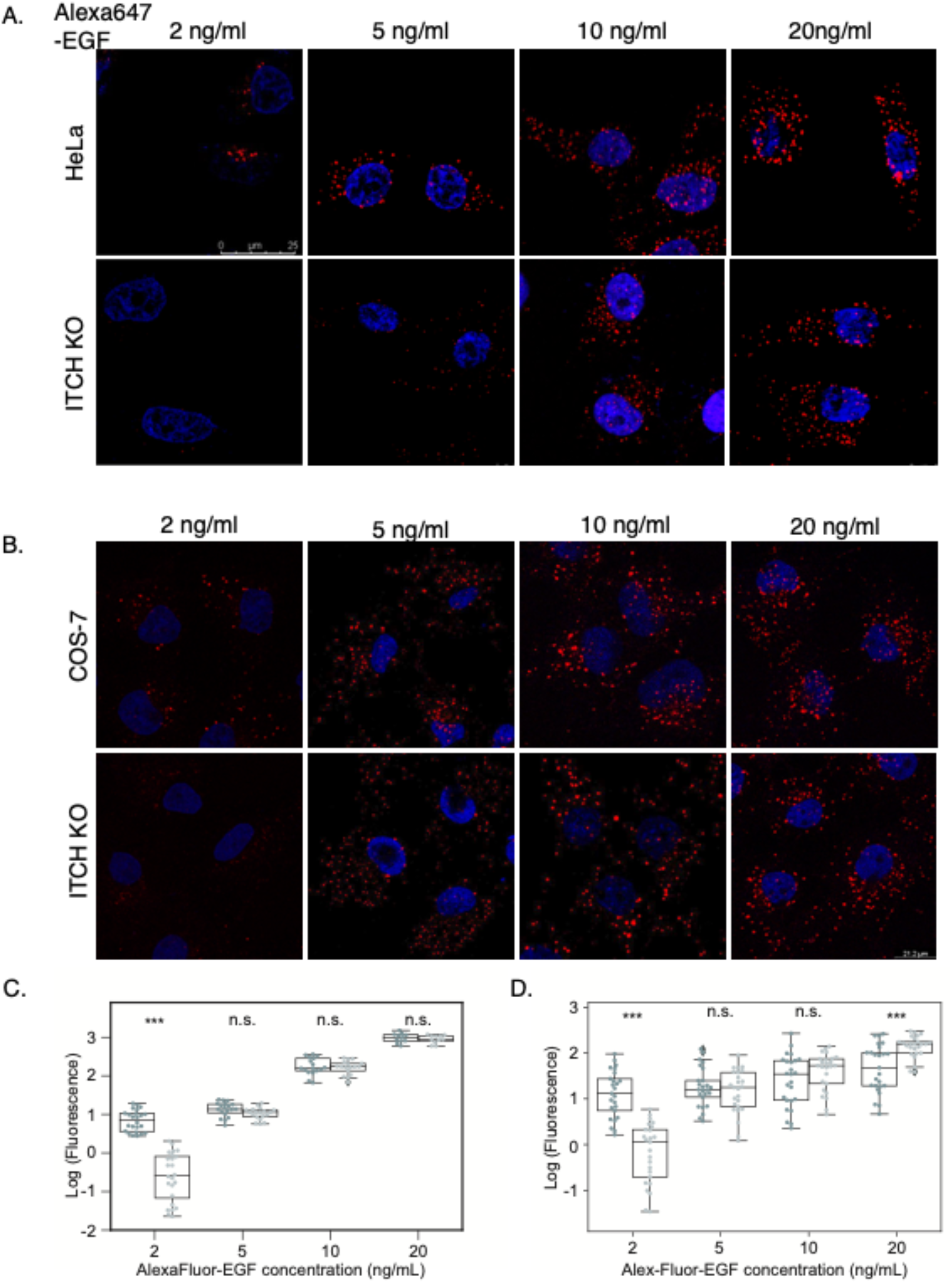
Endocytosis assays to detect clathrin-mediated internalization of EGF in HeLa and COS-7 ITCH^-/-^ cells. Uptake of AlexaFluor647-EGF at different concentrations in ITCH knockout A) HeLa cells and B) COS-7 cells versus the corresponding parental control cells. Cells were starved for 2 h and AlexaFluor647-EGF, diluted in serum-free media, was added at 20, 10, 5 and 2 ng/ml. Cells were gently acid washed to remove any extracellular bound EGF. Images were captured after 20 min of endocytosis. In all images, the nucleus is DAPI stained (blue). C) Quantification showing mean value of intracellular EGF fluorescence in HeLa. The reduced level of intracellular EGF in ITCH knockout cells compared to parent cells is only statistically significant at 2ng/ml. The difference is non-significant *(n.s)* at 5, 10 and 20ng/ml EGF. D) Quantification showing mean value of intracellular EGF fluorescence in COS-7. The difference is non-significant *(n.s)* at 5 and 10 ng/ml EGF. *t* test ***p<0.001. *n*=50 cells from one experiment, representative of three independent experiments. Scale bar, A) 25 μm and B) 21 μm.

### Alterations in EGFR CME are not due to a general endocytic defect

We next sought to determine if the deficit measured in EGFR internalization was due to a general impairment of CME in ITCH KO cells. We thus tested the effect of the absence of ITCH protein on the endocytosis of Transferrin. Unlike EGFR, the transferrin receptor is internalized in a constitutive and stimulation-independent manner. After 20 min of fluorescent transferrin uptake, there was no difference in the level of internalized fluorescent in ITCH KO cells compared to control (Fig. 2). Together, our data demonstrate that CME of EGFR is dependent on the ubiquitin ligase ITCH, although the CME machinery appears to function normally.

**Figure 2.**
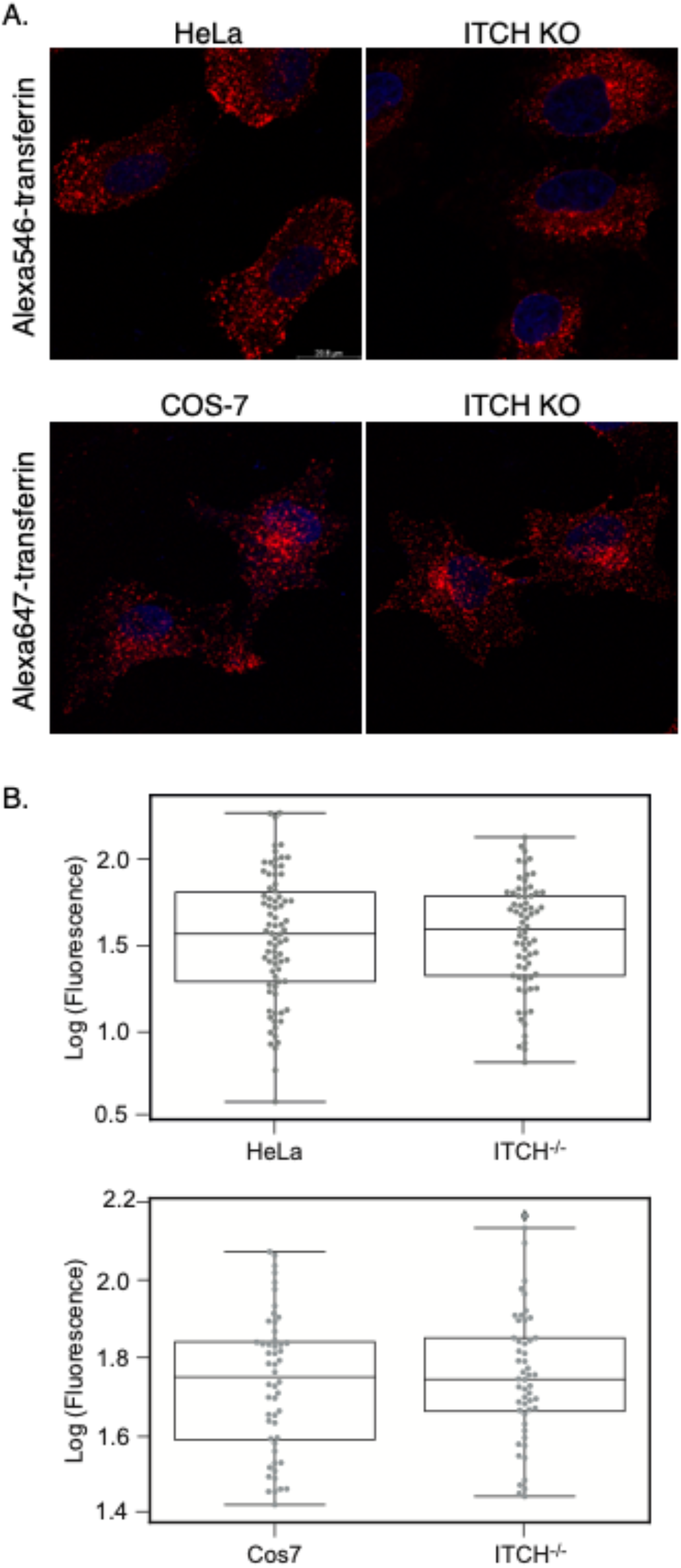
Endocytosis assays to detect clathrin-mediated internalization of transferrin in HeLa and COS-7 ITCH^-/-^ cells. A) HeLa and COS-7 ITCH^-/-^ cells vs parental cells allowed to internalize AlexaFluor-transferrin for 20min. Cells were gently acid washed to remove any extracellular bound transferrin. The nucleus is DAPI stained (blue). B) Quantification showing mean value of intracellular EGF fluorescence. The difference between ITCH knockout cells and parental cells is non-significant *(n.s)* in both cell lines. *t* test p>0.1, *n=80* cells (HeLa) and *n=50* cells (COS-7) from one experiment, representative of three independent experiments. Scale bar, 20 μm.

### EGFR degradation and signaling in ITCH KO cells

We next questioned if ITCH KO leads to any change in the rate of receptor degradation. For this purpose, we stimulated the cells at a low concentration of EGF and immunoblotted for total and phosphorylated EGFR. When compared to wild-type HeLa cells, the levels of total EGFR are comparable at all time points in ITCH KO cells (Fig. 3A). However, activated EGFR (pY1068 and pY1148) seemed to be slightly more stable in ITCH KO cells (Fig. 3B). Importantly, activated ERK1/2 remained activated longer in ITCH KO cells when compared to control (Fig. 3B). Together, these results suggest that EGFR signalling is moderately upregulated in ITCH knockouts.

**Figure 3.**
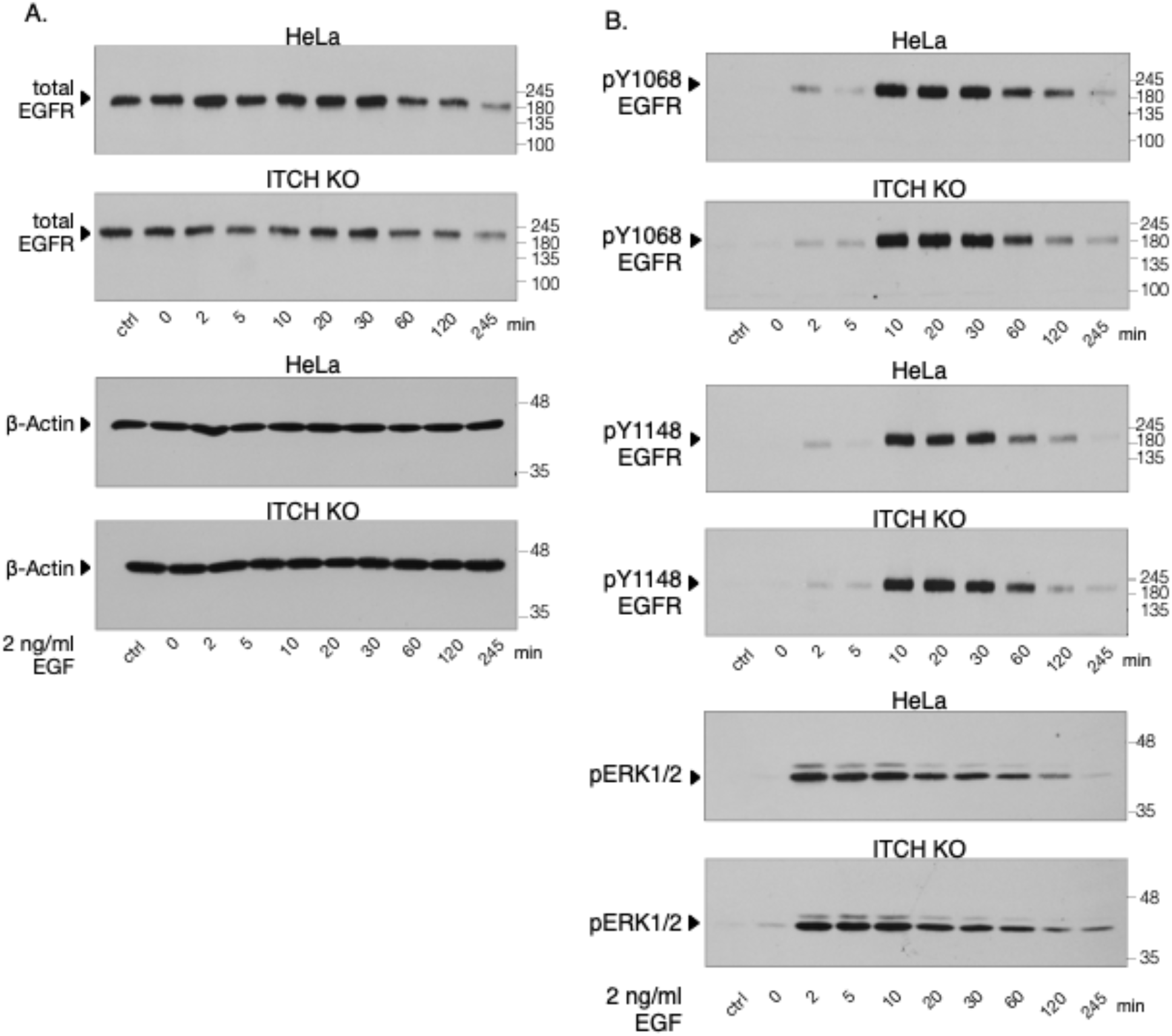
Degradation of activated EGFR upon CME in ITCH^-/-^ cells. After starvation for 2 h, WT and ITCH knockout HeLa cells were stimulated with 2 ng/ml recombinant EGF for the indicated amounts of time. Total lysates were subjected to immunoblotting with A) anti-total EGFR and β-Actin as a loading control and B) anti-pY1068 and pY1148 EGFR to follow the degradation the activated receptor over time and pERK1/2 to detect the effect on downstream signaling pathway. ITCH^-/-^ cells are showing a delay in phospho-EGFR degradation and prolonged ERK signaling.

### Mutations in the PRR of ITCH efficiently reduce SH3 domain binding

The ITCH PRR interacts with the SH3 domain of several endocytic proteins with the highest affinity for endophilin. The ITCH PRR is a 20 amino acid stretch (PSRPPRPSRPPPPTPRRP), bearing four canonical sites for recognition by SH3 domains; three class II sites and a single class I site (16). In previous studies, we determined that endophilin bound specifically to the class II motif RPPRPSR (residues 252–258) (9). We speculated that the arginines would be critical for the recognition of endophilin by full-length ITCH. We created four different point mutations in the PRR, replacing arginine residues with glutamic acid (Fig. 4A). We obtained R252E, R255E, R252,255,258E and R265,266E ITCH mutants. These GFP-tagged ITCH mutants were overexpressed in HEK-293T cells followed by a pull-down assay using 10 μg of the GST-SH3 domain of endophilin A2. While a single point mutation in R252 or R255 reduced the affinity of the SH3 domain for ITCH, mutating the R residues 252, 255 and 258 simultaneously nearly abolished the interaction between the SH3 of endophilin A2 and ITCH (Fig. 4B). Mutating residues 262-266 had little effect on the affinity between ITCH and endophilin A2-SH3 (Fig. 4B). To challenge the binding capacity of this triple mutant, we performed the same pull-down assay with increasing amounts of GST fusion proteins. Only negligible binding of Endophilin SH3 could be detected (Fig. 4C), while all other SH3 domains tested did not bind this mutant (Fig. S4).

**Figure 4.**
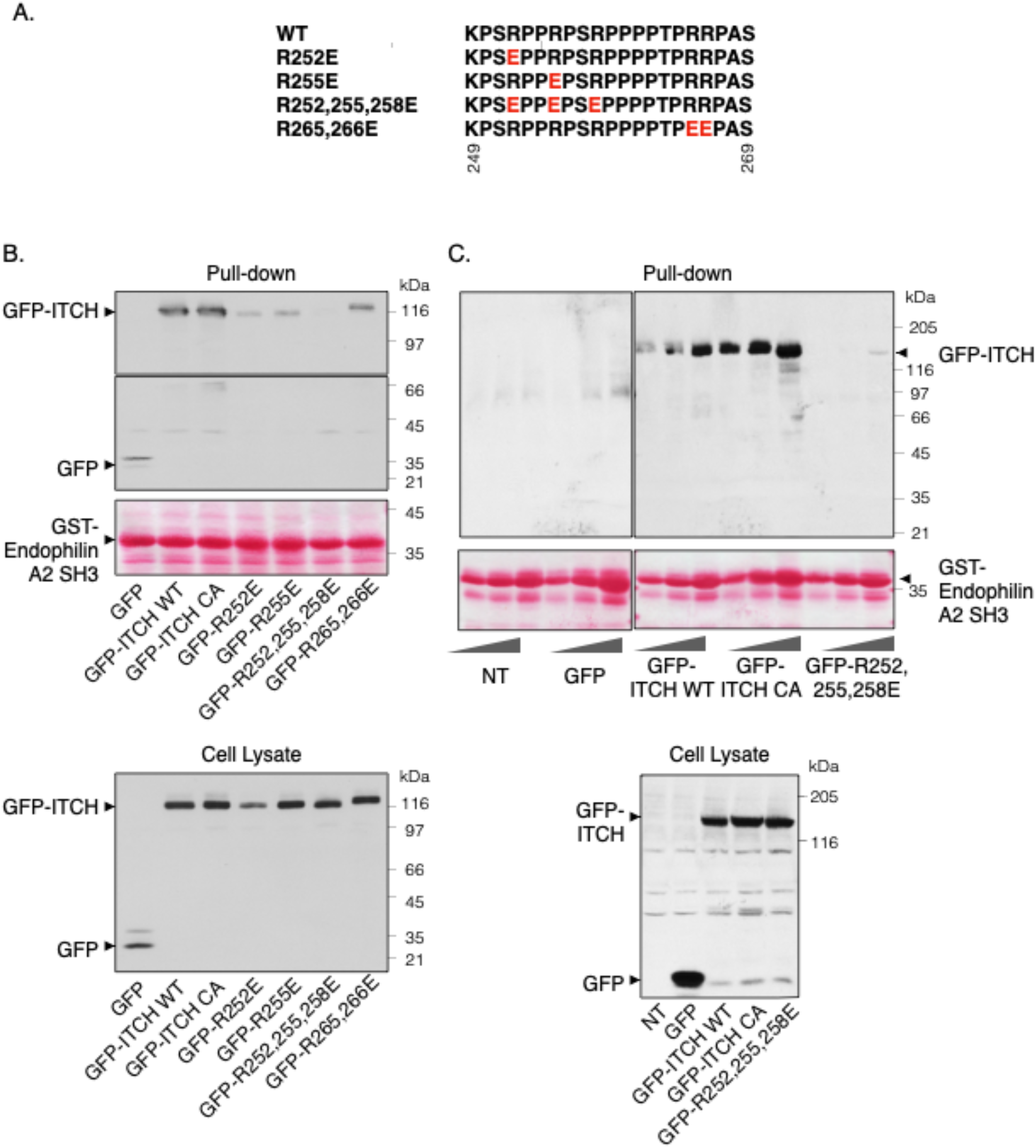
Pull-down assays testing the affinity of PRR mutant full ITCH to the SH3 domain of endophilin A2. A) Positions of the arginine residues replaced with glutamic acid in the region spanning from residue 249 to 269 in the PRR of ITCH. B) The cell lysate panel immunoblot shows the overexpressed levels of GFP, GFP-ITCH WT, ligand-dead-GFP-ITCH (CA), R252E, R255E, R252,255,258E and R265,266E in the total lysates of transfected HEK-293T cells. An aliquot of 10 μg of purified GST-endophilin A2 SH3 domain was incubated with lysates from these cells and the pull-down (top panel) shows GFP-tagged ITCH mutant proteins bound to endophilin A2 SH3. The triple arginine mutant does not bind to the SH3 domain of endophilin A2. C) A similar experiment focusing only on the R252,255,258E mutant in comparison to GFP-ITCH WT and CA in the presence of a gradient of fusion protein concentration (5, 10 and 20 μg). The cell lysate panel shows anti-GFP western blot of the starting material. The anti-GFP western blot of the pulldown shows GFP-tagged ITCH proteins bound to the different concentrations of purified fusion protein. ITCH residually binds to 20 μg of endophilin A2 SH3. At the bottom of each pull-down experiment is the Ponceau-S staining for the fusion protein.

### Rescue of EGFR internalization

In order to further demonstrate the contribution of ITCH to CME of EGFR, we performed a rescue experiment using GFP-ITCH wild-type, catalytically inactive (CA) and SH3-binding deficient (R252,255,258) mutants. While EGFR internalization was re-established by overexpression of GFP-ITCH wild-type, both mutants failed to rescue the phenotype (Fig. 5). This indicates that both ITCH ubiquitin ligase activity and scaffolding capacities are needed for CME of EGFR.

**Figure 5.**
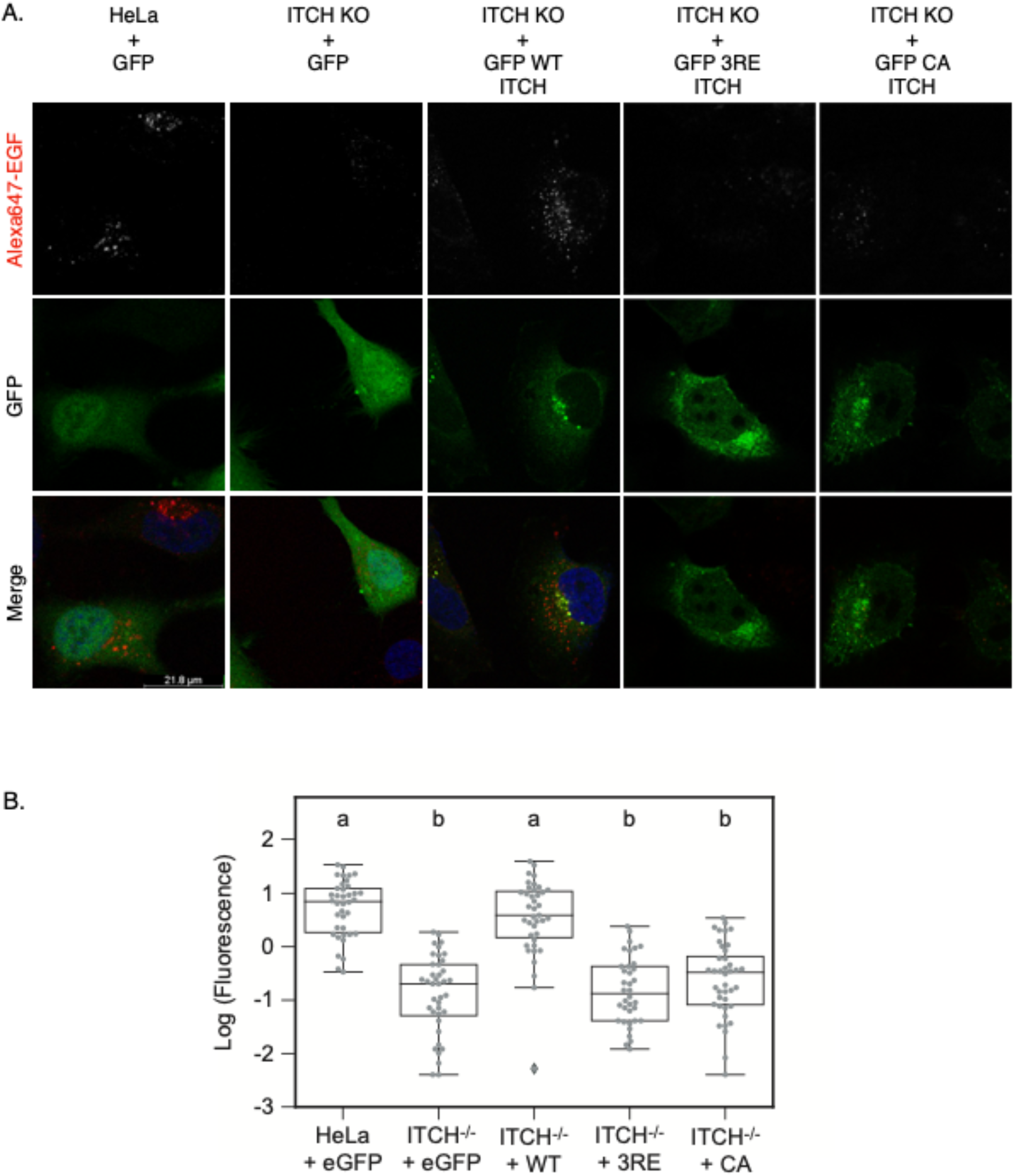
Rescue experiment using ITCH CA and triple arginine mutant. A) As a control (first lane), WT and ITCH Knockout HeLa cells were overexpressed with GFP. ITCH knockout cells expressing GFP-ITCH WT, ligase-dead (CA) and R252,255,258E (3RE) were allowed to internalize 2 ng/ml AlexaFluor647-EGF (red) for 20min. Top panel is shown in gray scale with increased luminosity for clarity. The nucleus is DAPI stained (blue). B) Quantification of the mean value of intracellular EGF fluorescence and statistical significance determined by *Anova* and *Bonferroni* tests. a and b refer to two different groups of values significantly different one from another. Parent HeLa cells and ITCH^-/-^ cells overexpressed with GFP-ITCH WT (group a) are significantly different from ITCH^-/-^ cells overexpressed with GFP, GFP-ITCH CA or R252,255,258E (group b). *n=40* cells from one experiment, representative of three independent experiments. Scale bar, 21 μm.

## Discussion

EGFR uptake at the plasma membrane controls many cellular functions including proliferation and migration. This process is highly regulated, and a dysregulation of EGFR trafficking is strongly associated with oncogenesis (20). Here we focus on the role played by the ubiquitin ligase ITCH to control the internalization and signaling of EGFR. ITCH binds to c-CBL through its WW domains, and its PRR region interacts with Endophilin, two key players in EGFR endocytosis (7,8).

When using low concentrations of EGF, activated EGFR uses only the clathrin-mediated route for internalization (6). We found here that the absence of ITCH caused a deficiency in the internalization of low levels of EGF, implicating CME. Previous studies have implicated ITCH in the regulation of CME. For example, ITCH can bind many SH3 domain-containing proteins including intersectin-1, amphiphysin, PACSIN and β-PIX (9). Endophilin is recruited to the endocytic site through CIN85/CBL and is a major binding partner to ITCH (9,16). Our results reveal for the first time that the CME of stimulated EGFR is dependent on ITCH.

The CBL family of ubiquitin ligases are known to ubiquitinate EGFR post-activation and send it to degradation in lysosomes. CBL was shown to be a multi-adaptor protein that recruits CIN85 and endophilin to form a complex with activated EGFR (13). CBL is polyubiquitinated by ITCH in a degradative way, which suggests that ITCH is a negative regulator of EGFR degradation (7,21). Knockdown of CBL in HEK-293T cells did not significantly affect the level of EGF-activated EGFR internalization at 100 ng/ml but delayed its ubiquitination and degradation (22). In porcine aortic endothelial cells, CBL knockdown reduces the level of internalized receptor up to 40-50% compared to the control, showing that this internalization is dependent on CBL (23).

Sorkin et al. later identified ubiquitination as part of an essential mechanism through which EGFR can be internalized, when showing that mutating 21 lysine residues in the tyrosine domain of EGFR significantly reduced the receptor internalization (3). This clathrin-mediated internalization also depends on adaptor proteins like the BAR and SH3 domain-bearing protein, endophilin. At synapses, an endophilin triple knockout impaired uncoating and provoked an accumulation of clathrin-coated vesicles, delaying endocytosis and impairing synaptic transmission (12). In breast cancer cells, an EGF internalization deficiency is noticeable after the silencing of endophilin A2 (24).

In rescue experiments, the triple arginine mutant failed to recover the normal level of intracellular EGF, indicating that ITCH PRR availability is important to complete the endocytic complexes. While it is not rare for a ubiquitin ligase to play the role of an adaptor protein, it is uncommon for one to bear a PRR. Therefore, ITCH could participate in EGFR endocytosis through the recruitment of endophilin and/or other endocytic proteins. PRR/SH3 interactions are known to drive the formation of CCPs. For example, dynamin, the GTPase responsible for the fission of CCPs, uses its PRR to bind the SH3 domains of endophilin and amphiphysin (25,26). In addition, these interactions can control the endocytic trafficking and degradation of the cargo. The Rin1/STAM complex functions as a regulator of EGFR trafficking. The Rab5 exchange factor Rin1 (Ras interaction/interference 1) binds the SH3 domain of STAM (signal transducing adaptor molecule) via its PRR. Both proteins colocalize to the stimulated EGFR at endosomes, to guide the sorting of the receptor to lysosomes (27). A similar colocalization between ITCH and endophilin A1 is seen after overexpression of both proteins and EGF stimulating the cells (8). The SH3 domain-containing protein intersectin1 is also another substrate of ITCH acting at the endocytic site. Intersectin1 stimulates the ligase CBL, thus stimulating EGFR ubiquitination. By binding the PRR of both CBL and its inhibitor sprouty 2, intersectin disrupts the sprouty 2-CBL interaction, leading to an enhanced EGFR signaling (28). The isoform 1s of intersectin is directly monoubiquitinated by ITCH, a modification that does not lead to its degradation (29).

Similarly, our data show that an ITCH ligase dead mutant is unable to restore a normal level of internalized EGF. As a substrate of ITCH at the endocytic vesicle, CBL ubiquitination can be directly affected by the loss of a catalytically active ubiquitin ligase. Any change in the ubiquitination levels of ITCH substrates at the plasma membrane can disturb the formation of the protein complexes carrying the cargo. The ubiquitination of the adaptor protein eps15 gives an example on the importance of this post-translational modification in completing a normal endocytic complex. Eps15 ubiquitination by Parkin prevents it from binding ubiquitin on the receptor, a modification that delays the internalization and degradation of EGFR, causing an amplified PI3K-Akt signaling (30).

Transferrin endocytosis is independent of its ligand binding. When followed in ITCH^-/-^ cells, there is no difference with control cell lines. This demonstrates that ITCH recruitment and implication in endocytosis is dependent on receptor activation. ITCH interacts with CBL and Grb2 (31), two proteins specifically recruited by activated EGFR.

In the absence of ITCH, the endogenous levels of total EGFR, CBL and endophilin A2 remain unchanged between the parental and ITCH^-/-^ HeLa at any given time. However, after stimulation with low EGF levels, the phosphorylated EGFR follows a slower degradation rate in ITCH^-/-^ compared to WT cells, accompanied by a prolonged stimulation of pERK. This prolonged MAPK activation correlates with cancer progression (32). In the future, the ubiquitination state of the stimulated EGFR and c-CBL should be assessed in the ITCH^-/-^ cell line in order to better understand any events that could be related to the internalization deficiency. In order to confirm if the regulatory effect of ITCH is specific to EGFR endocytosis, another ligand-dependent receptor internalization should be studied in the ITCH^-/-^ cell line. c-Met is another receptor known to use the endophilin-CIN85/CBL during endocytosis, which makes it a good candidate for endocytosis testing in the ITCH KO cell line (33). CXCR4 (34) and Notch (35) are also receptors ubiquitinated by ITCH, a modification that affects their trafficking and degradation,

In summary, we discover that clathrin-mediated endocytosis of EGFR is dependent on the ubiquitin ligase ITCH. We also determine that ITCH does not only function as a ubiquitin ligase but also as an adaptor protein to facilitate this process.

## Supporting Information

**Figure S1.**
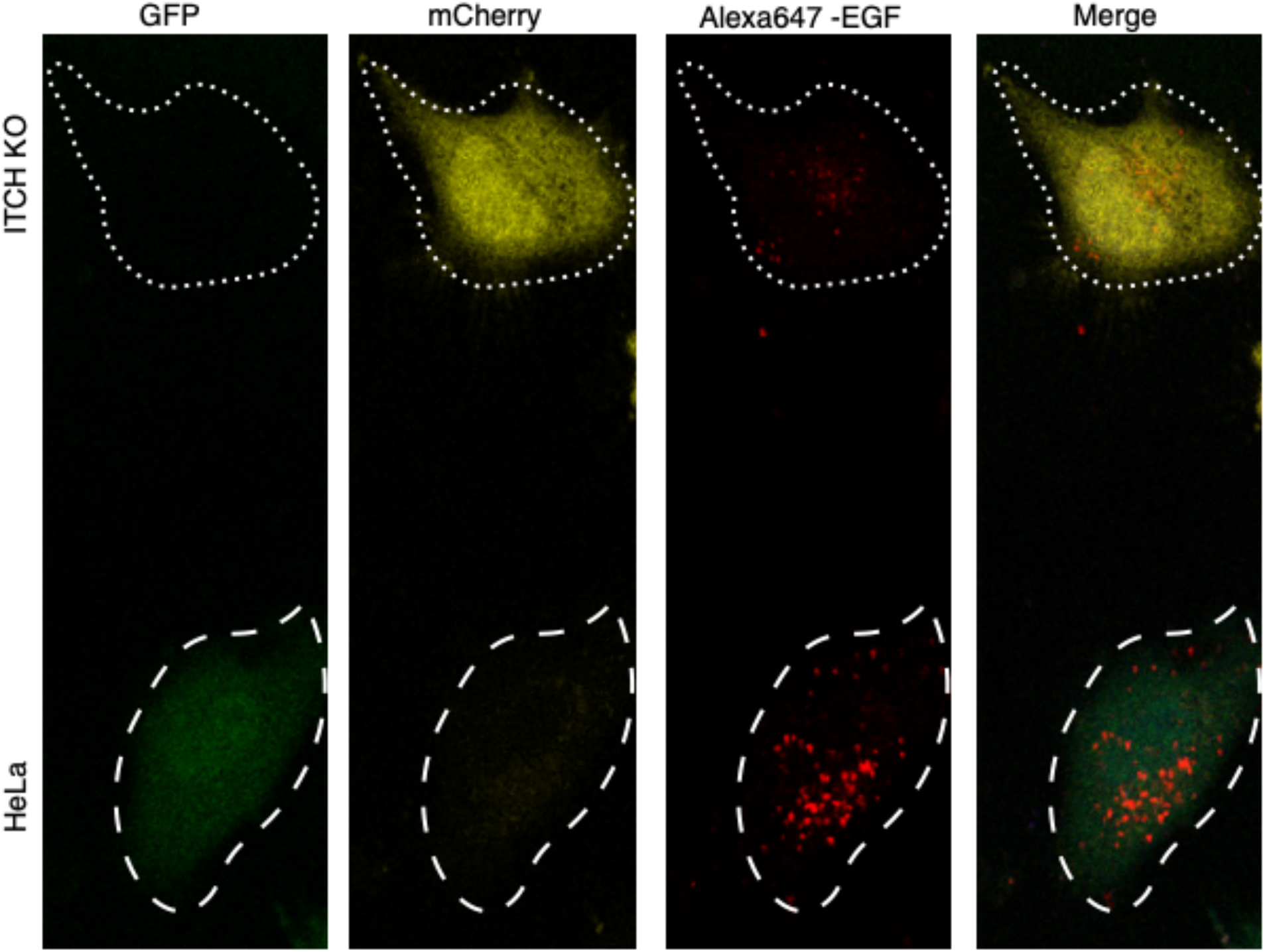
Endocytosis assay to detect clathrin-mediated internalization of EGF by a mosaïque of parent and ITCH^-/-^ cells. HeLa cells are overexpressing GFP (green) and ITCH^-/-^ cells mCherry (yellow). Cells were captured in the same microscopic field after 20min of endocytosis using 2ng/ml of AlexaFluor-647 EGF. HeLa cell is shown with loosely dashed lines, ITCH^-/-^ cell with dash dotted lines.

**Figure S2.**
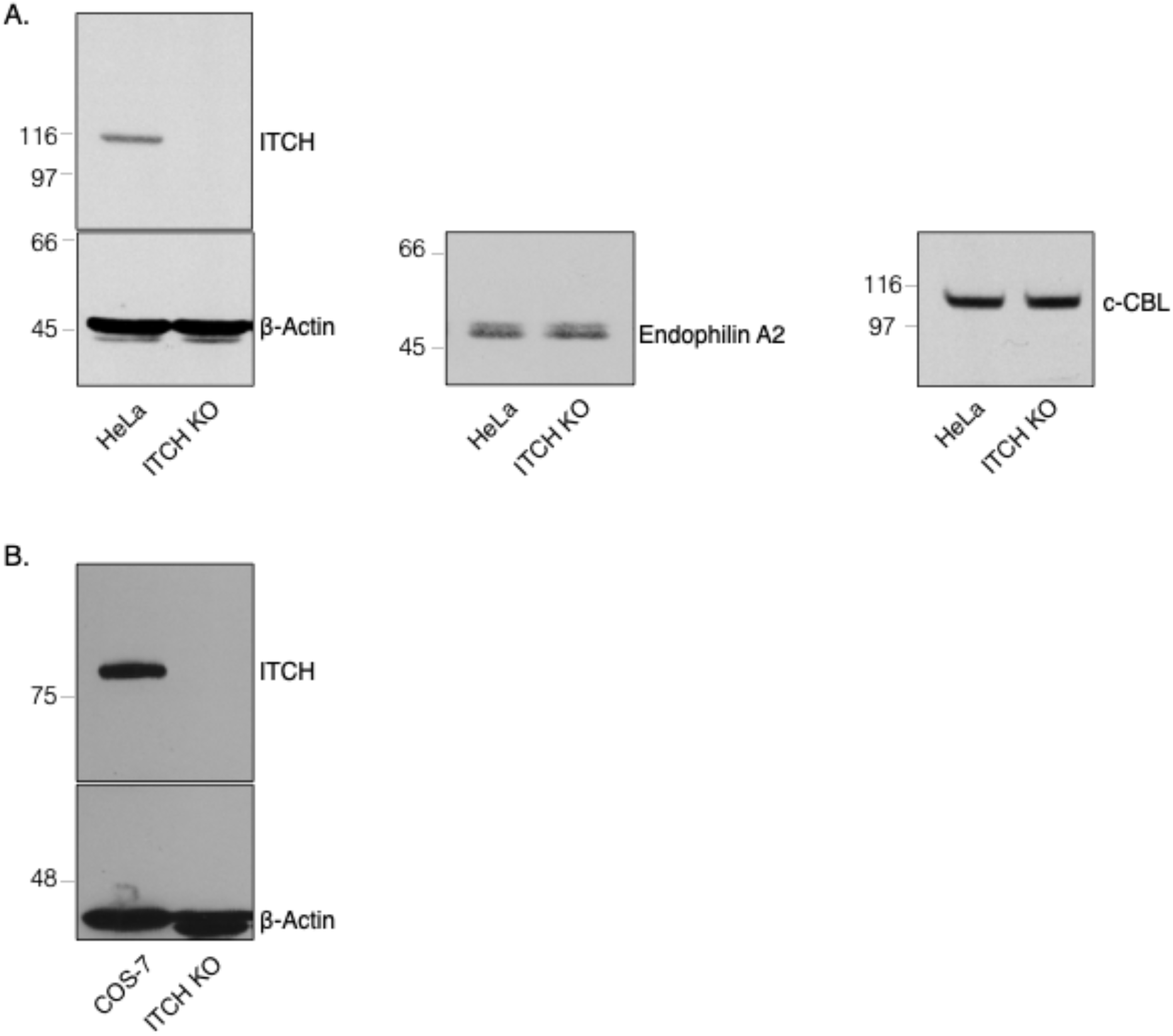
A) Western blot for the determination of the endogenous levels of ITCH partners involved in CME of EGFR. The cells at steady-state were lysed, and total lysates were subjected to SDS-PAGE and immunoblotting with antibodies against the proteins shown. B) Western Blot using anti-ITCH to validate the knockout in COS-7 cells.

**Figure S3.**
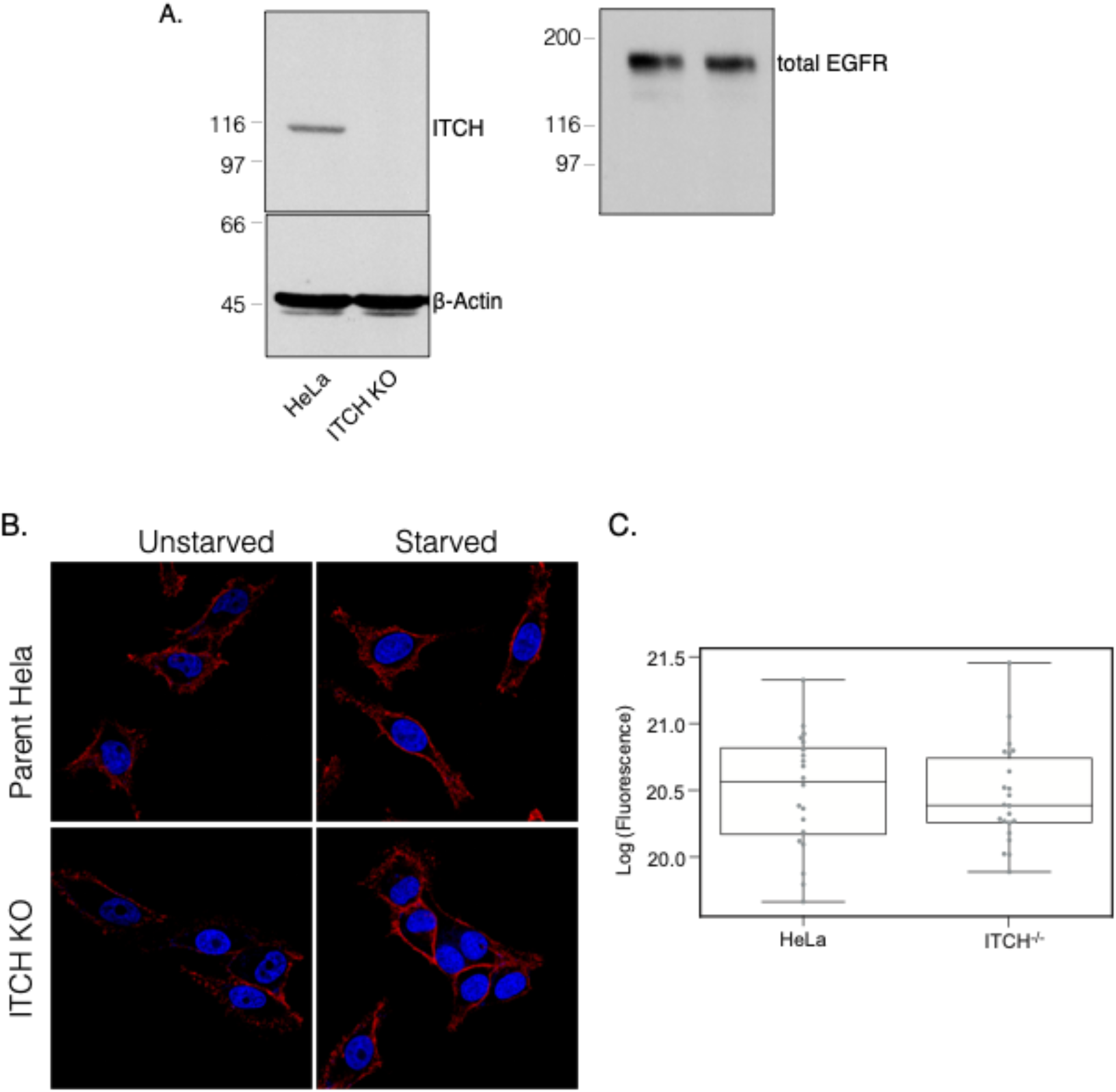
Total levels of EGFR compared in HeLa and ITCH^-/-^ cells by Western-Blot and Immunofluorescence. A) Western blot for the determination of the endogenous total levels of EGFR. The cells at steadystate were lysed, and total lysates were subjected to SDS-PAGE and immunoblotting. B) To determine the level of extracellular EGFR, an antibody that recognizes the extracellular domain of EGFR was used, followed by Fluor568-conjuguated secondary antibody. Images of starved or unstarved cells were captured at steady state. The red signal shows non internalized EGFR. The nucleus is DAPI stained (blue). C) Quantification showing mean value of extracellular EGF fluorescence in unstarved cells. The difference is non-significant *(n.s)* based on *t* test, p>0.1, *n=20* cells from one experiment.

**Figure S4.**
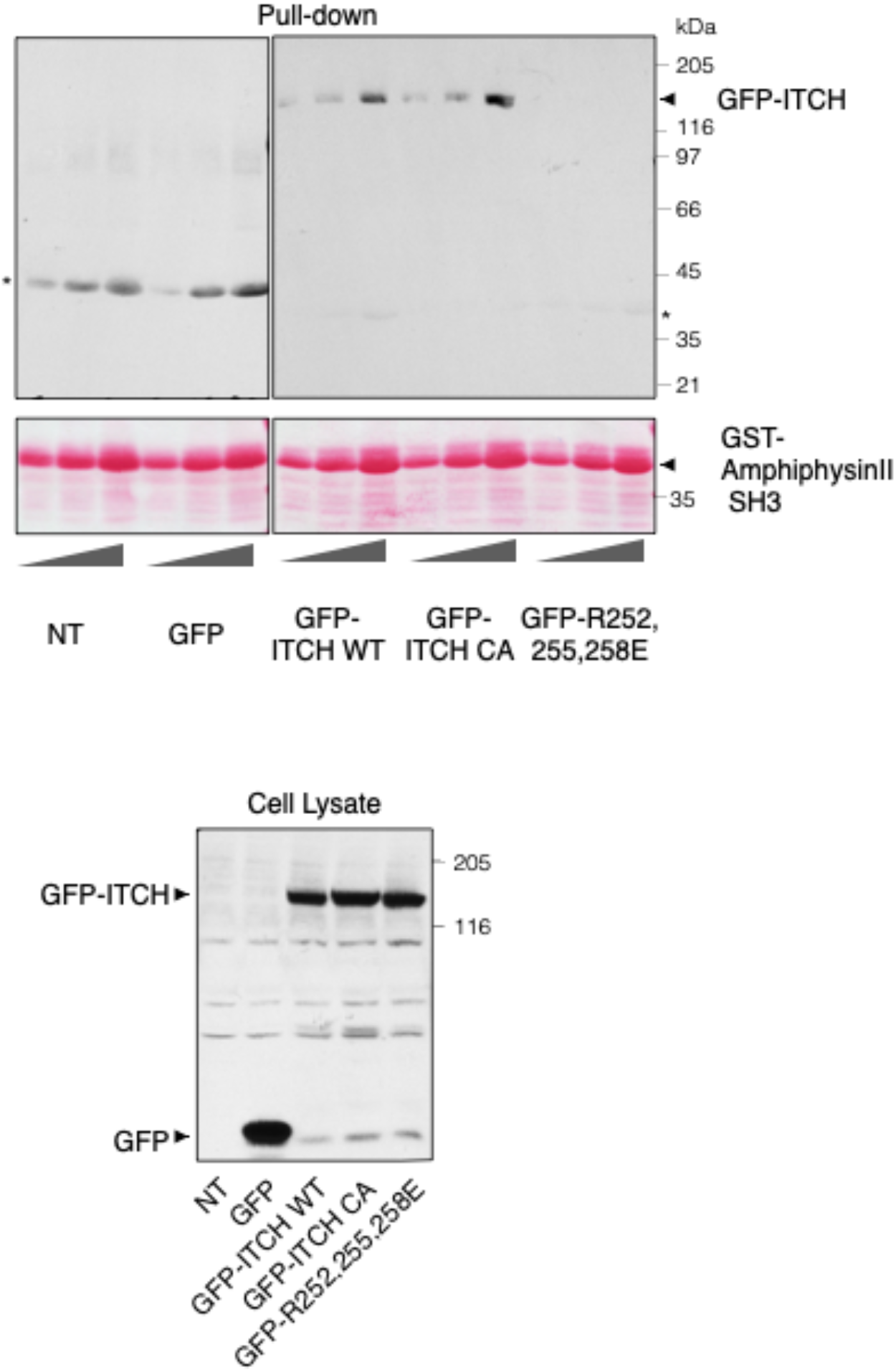
Pull-down assays testing the affinity of PRR mutated full ITCH to the SH3 domain of amphiphysin II. The cell lysate panel immunoblot shows the overexpressed levels of GFP, GFP-ITCH WT, CA and R252,255,258E in the total lysates of transfected HEK293T cells. A gradient of purified GST-amphiphysin II SH3 concentrations (5, 10 and 20 μg) was incubated with lysates from these cells and the pull-down shows GFP-tagged ITCH proteins bound to amphiphysin II SH3. At all concentrations, the triple arginine mutant does not bind to the SH3 domain of amphiphysin II.

